# Structural diversification of phage tail fibres enables recognition of diverse type IV pili

**DOI:** 10.64898/2026.03.17.712343

**Authors:** Ikram Qaderi, Hanjeong Harvey, Yao Shen, Ylan Nguyen, Amogelang R. Raphenya, Isabelle Chan, Alba Guarné, Andrew G. McArthur, Lori L. Burrows

**Author notes:** For correspondence: Dr. Lori L. Burrows.

## Abstract

Viruses must recognize receptors on host surfaces to initiate infection, but these receptors can evolve rapidly, posing a fundamental challenge to viral persistence. The type IV pilus of *Pseudomonas aeruginosa* is an ideal model system to study virus-receptor coevolution because its major pilin subunit PilA exhibits extensive sequence and chemical diversity while remaining essential for phage attachment. Here, we combined large-scale comparative genomics with structural and functional analyses to determine how pilus-dependent phages maintain infectivity despite extensive receptor diversification. Pilin variation was concentrated at solvent-exposed regions, altering filament surface chemistry while preserving key subunit-subunit interfaces. Despite this variation, phages recognized divergent pilins more effectively than polyclonal antisera. However, phages differed markedly in their sensitivity to receptor perturbation: some required electrostatic and structural compatibility, whereas others tolerated substantial receptor modification, including the presence of post-translational glycosylation. Comparisons of AlphaFold3 structural models revealed two distinct tail fibre architectures associated with those phenotypes. Phages encoding tail fibres with conserved receptor-binding domains were more sensitive to receptor perturbation within a given pilin background, while those with structurally diversified binding regions infected strains expressing highly divergent pilins. Together, these findings suggest that modular diversification of receptor-binding proteins provides a structural route by which phages accommodate receptor evolution.

**SIGNIFICANCE:** Understanding the determinants of viral host range is essential for explaining how viruses persist in the face of receptor diversification. Using pilus-dependent bacteriophages that bind diverse type IV pilins in *Pseudomonas aeruginosa*, we show that differences in tail fibre architecture are associated with distinct sensitivities to receptor variation. Phages with structurally conserved receptor-binding regions are sensitive to receptor perturbation, whereas those with diversified binding architectures remain infectious across strains expressing highly divergent pilins. Our results identify a structural correlate of receptor recognition breadth and reveal how receptor-binding protein architecture contributes to viral host range and adaptation. These data can inform the selection or engineering of broad-host range phages for therapeutic applications.

## INTRODUCTION

Surface structures are among the most evolutionarily dynamic elements in bacteria. They mediate interactions with the environment, hosts, competing microbes, and viruses, and are therefore subject to multiple selective pressures (1, 2). For bacteriophages (phages), receptor binding is the critical first step of infection, yet many receptors exhibit substantial sequence and structural variation across host populations (3, 4). How phages maintain infectivity despite receptor diversification remains a central question in virus-host coevolution.

Type IV pili (T4P) provide a powerful system to investigate this problem. These dynamic, retractile filaments are assembled from thousands of pilin subunits and are broadly distributed in both archaea and bacteria (5, 6). In *Pseudomonas aeruginosa*, T4P are important virulence factors that mediate motility, surface attachment, microcolony formation (7, 8), and serve as receptors for diverse phages (9–11). Imaging studies show that phages from various families can attach along the length of the pilus or at its tip and are brought to the cell surface by pilus retraction (12–15), suggesting that different phages engage the same receptor using distinct attachment strategies.

New mechanistic insight into pilus recognition has emerged recently, primarily from work on tailless single-stranded RNA phages such as PP7, which bind defined regions of the major pilin PilA and are highly sensitive to single amino acid substitutions (14, 16, 17). However, most characterized *Pseudomonas* phages are tailed double-stranded DNA viruses (18), and how they engage structurally variable pilins remains poorly defined. Structural and genetic studies of dsDNA phages have proposed multiple attachment strategies, including baseplate-mediated filament binding or recognition of minor pilins located at the pilus tip (15, 19). These interactions are mediated by tail fibres, which are among the most structurally and evolutionarily diverse components encoded by phage genomes and play a central role in determining host range (20–22). Structural diversity among tail fibres suggests differences in their pilin-binding interfaces, which may help explain variation in receptor specificity and tolerance. However, direct structural information for tail fibre-pilus complexes remains limited.

In *P. aeruginosa*, the major pilin PilA exhibits extensive sequence diversity across strains (23, 24). Population-level surveys identified five major pilin groups, defined in part by accessory genes encoded adjacent to *pilA* and by variation in length of the C-terminal disulfide-bonded loop (24). The accessory proteins can post-translationally modify some PilA variants (25–27), which alters susceptibility to a subset of phages (28). Structural and genetic variation in PilA therefore provides a natural system to study how receptor diversification influences phage infection. However, to what extent phages can tolerate pilin diversity and whether this tolerance is predictably encoded by tail architecture remain unclear.

We hypothesized that pilus-dependent phages use distinct modes of receptor engagement that differ in their tolerance to pilin variation, and that these differences are associated with tail module architecture. To test this, we integrated population genomics, structural analysis and modelling, targeted mutagenesis, and comparative phage phenotyping. We defined the distribution of PilA variants across more than 1,300 *P. aeruginosa* genomes, mapped sequence variation onto pilin structures and models, assessed antigenic and phage recognition using a recombinant pilin library, and examined whether tail module architecture segregates with sensitivity to receptor mutation and modification.

Here we show that pilus-dependent phages exhibit two distinct patterns of tolerance to pilin variation that align with tail module architecture. Phages encoding structurally conserved tail fibres show narrow recognition breadth and are sensitive to electrostatic perturbations at the solvent-exposed pilus surface, whereas phages encoding structurally diversified tail fibres tolerate substantial pilin sequence variation and post-translational modification. These findings suggest tail module architecture contributes to differences in receptor recognition breadth and provide a structural and evolutionary framework for understanding how phages accommodate receptor diversification.

## RESULTS

### PilA diversity is concentrated at solvent-exposed regions of the pilus filament

To define the receptor diversity encountered by pilus-dependent phages, we analyzed PilA sequences from 1,365 nonredundant *P. aeruginosa* genomes and mapped them to their multilocus sequence types (MLST) (29). This approach reduced potential clonality and revealed which pilins were most widely distributed within the species. We identified 53 different PilA sequences corresponding to 319 unique pilin-MLST combinations, all of which segregated into the five previously defined pilin groups (**Figure 1A,B**; **Supplementary File F1**) (24). The majority of the 319 unique pilin-MLST combinations had group I (∼46%) or group II (∼36%) pilin genes, a distribution consistent with our prior survey (24), with the rest having groups III (∼9%), IV (∼3%), or V (∼6%) (**Figure 1B**).

**Figure 1.**
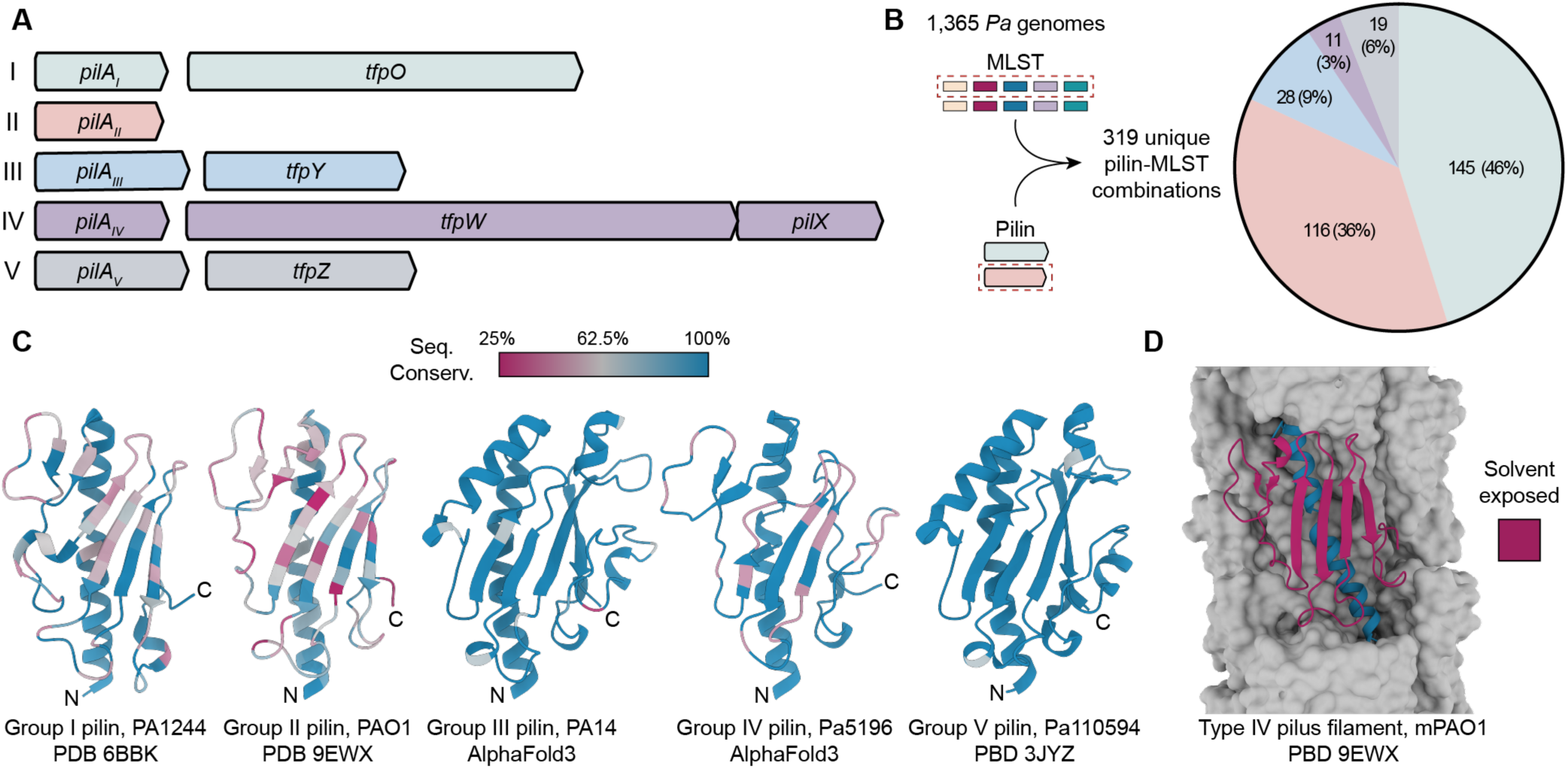
PilA diversity is concentrated at solvent-exposed regions of the pilus filament. **A.** Genetic organization of the five pilin groups in *P. aeruginosa*, defined by the pilin structural gene (*pilA*) and associated accessory genes required for pilin modification or assembly (24). **B.** Distribution of pilin groups among 319 unique pilin-MLST combinations identified from 1,365 nonredundant *P. aeruginosa* (*Pa*) genomes. Group I and group II pilins predominate, with smaller fractions belonging to groups III, IV, and V. **C.** Mapping of sequence conservation onto representative pilin structures and AlphaFold3 (32) models. Experimentally determined structures are shown for group I (this study), group II (38), and group V (47) pilins, and AlphaFold3 models are shown for group III and IV pilins. Residues are coloured according to sequence conservation within pilin groups, with conserved residues in blue and variable residues in red. **D.** Type IV pilus filament model showing sequence divergence is concentrated at solvent-exposed regions (red) of the globular domain, whereas structurally conserved regions, including the N-terminal α-helix and buried core residues, remain highly conserved. AlphaFold3 models coloured by pLDDT values are shown in **Supplementary Figure S2B**.

Sequence divergence varied substantially between groups. Group II pilins displayed the greatest within-group sequence divergence (∼59% overall similarity), whereas pilins in groups I and IV were more conserved (>81%), and those within groups III and V were nearly identical (>97%) (**Supplementary Table S1**). Among group II pilins, phylogenetic analyses identified two new subgroups, designated IIB and IIC, that were ∼33-38% similar to the group II pilins described previously, now renamed IIA (**Supplementary Figure S1)**. Despite their sequence divergence, IIB and IIC pilins retain the defining features of group II, including 12 residues between the conserved cysteines delimiting the C-terminal disulfide-bonded loop and the absence of accessory proteins (24).

To examine how this sequence variation is distributed structurally, we solved a 1.7 Å crystal structure of PilA^1244^, a well-studied group I pilin (30, 31) and one of the most prevalent variants in our dataset (**Supplementary Figure S2A**; **Supplementary Table S2**). PilA^1244^ adopts a canonical type IV pilin fold, consisting of a long N-terminal α-helix connected to a 4-stranded anti-parallel β-sheet by an αβ loop. The C-terminus of the protein contains a 17-residue disulfide-bonded loop that pins it to the last β strand, and terminates in a Ser that serves as the site of post-translational modification with an LPS O-antigen unit (30, 31). Mapping within-group sequence conservation onto this structure, as well as onto representative structures or AlphaFold3 (32) models for the other four pilin groups, revealed that sequence divergence is concentrated almost exclusively at solvent-exposed residues within the globular domain, whereas buried residues are highly conserved (**Figure 1C, D**). These data indicate that pilus-dependent phages encounter substantial variation in pilin surface chemistry at solvent-exposed faces of assembled filaments.

### Phages are more tolerant of pilin sequence divergence than polyclonal antisera

Diversification of solvent-exposed residues is expected to alter recognition by antibodies and phages, therefore pilin variation in *P. aeruginosa* likely reflects selection imposed by both immune recognition and phage predation (23). However, whether sequence changes that preclude antibody binding also impair receptor recognition for pilus-dependent phages was unclear. If phages recognize precise molecular epitopes, similar to antibodies, pilin diversification should limit both antibody recognition and phage infection.

To isolate the specific effects of pilin sequence variation on receptor recognition, we constructed a defined library of representative pilins expressed in a common mPAO1 *pilA* mutant background. Pilins were selected based on their prevalence among the 319 unique pilin-MLST combinations identified in our dataset (**Figure 1B**, **Supplementary File F1**). Expression in a single genetic background eliminates confounding host-specific phage resistance mechanisms, allowing receptor compatibility to be assessed independently of downstream defence systems. All recombinant strains produced functional pili, as demonstrated by arabinose-inducible restoration of twitching motility (**Supplementary Figure S3A**), and surface piliation was confirmed by SDS-PAGE analysis of sheared pili (**Supplementary Figure S3B**).

We first asked whether pilin sequence divergence alters antigenic recognition. Whole cell lysates were probed with polyclonal rabbit antisera raised against representative pilins from each group. For groups I (this study), II (33), and IV (27), antisera were generated against N-terminally truncated recombinant pilins expressed in *Escherichia coli*, whereas antisera for groups III and V (34) were raised against pili sheared from *P. aeruginosa*. In both cases, immunogens primarily exposed the soluble globular domain of PilA and unlike monoclonals, we expected them to recognize a mixture of epitopes. Each antiserum reacted strongly with its cognate antigen but showed little or no cross-reactivity with divergent pilins, consistent with their limited sequence identity (**Figure 2A, Supplementary Table S1**). There were some instances of unexpected but weak cross-reactivity. For example, the group I pilin was weakly recognized by anti-group IIA antiserum, which failed to recognize IIB or IIC pilins (**Figure 2A**). Conversely, the group I antiserum failed to recognize the group II pilins (**Figure 2A)**. These data suggest that pilin diversification can block antigenic recognition, consistent with previous studies showing that vaccination with pilins protects only against homologous challenge (35–37).

**Figure 2.**
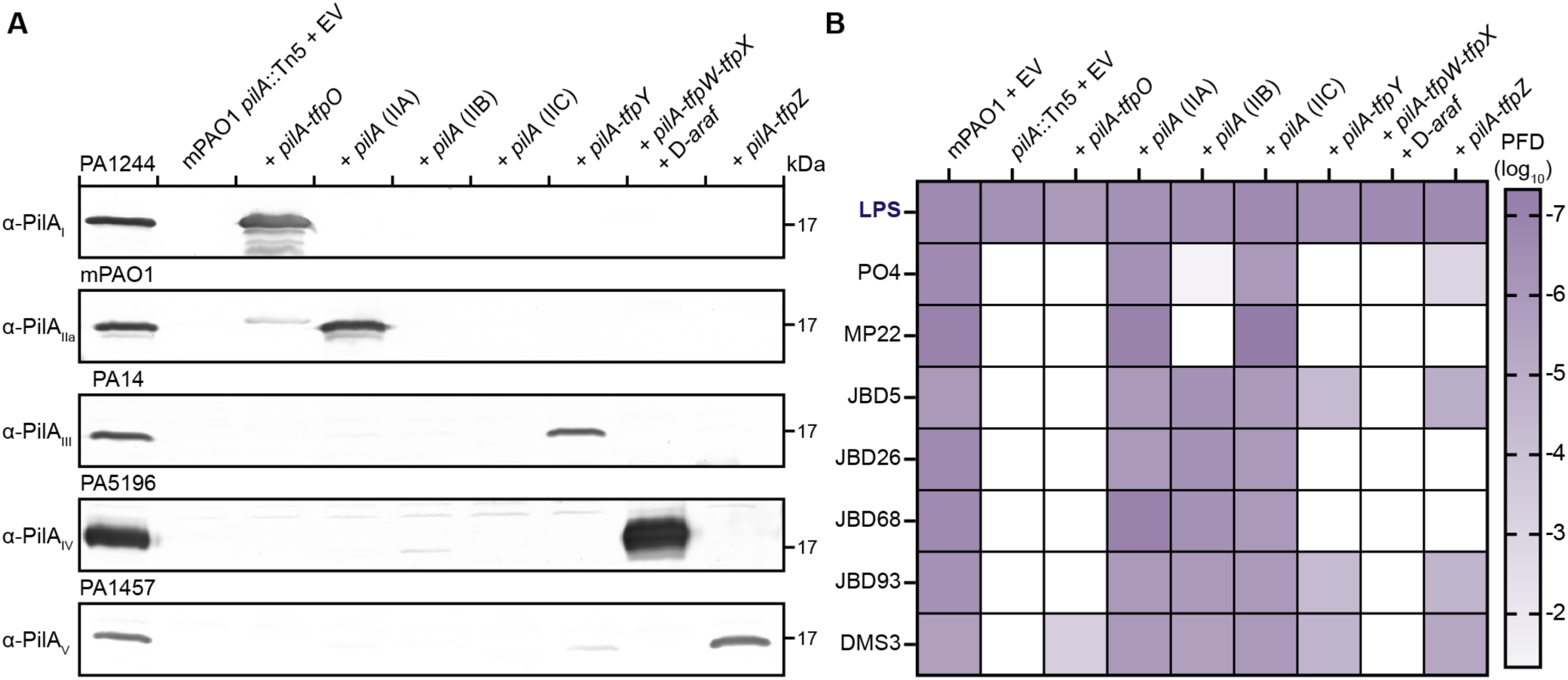
Phages are more tolerant of pilin sequence divergence than polyclonal sera. **A.** Immunoblot analysis of whole-cell lysates expressing representative pilins probed with group-specific rabbit polyclonal antisera. Except where indicated, pilin alleles were expressed from plasmids in a common mPAO1 *pilA* mutant background together with their associated accessory proteins required for pilin assembly or modification. For group IV pilins, genes required for D-arabinofuranose (D-*araf*) biosynthesis and transfer were coexpressed to enable pilin glycosylation. The first lane in each blot shows the native pilin expressed from its endogenous strain background, serving as a positive control for each antiserum. Antisera display strong reactivity toward cognate pilins but little to no cross-reactivity between pilin groups. Blots are representative of three independent experiments. **B**. Phage infectivity across the same pilin library expressed in the mPAO1 *pilA* mutant background. Susceptibility to pilus-dependent phages (PO4, MP22, JBD5, JBD26, JBD68, JBD93, and DMS3) and the LPS-targeting control phage Kipling (‘LPS’) is presented as plaque-forming dilution (PFD; log_10_ scale), defined as the highest dilution producing detectable plaques. Plaque assays used to generate heat map can be found in **Supplementary Figure S4**.

We next asked whether the same sequence differences that prevent antibody recognition would impair phage receptor recognition. Pilin library strains were infected with seven different pilus-dependent phages and a control lipopolysaccharide (LPS)-targeting phage, Kipling.

Because all strains share an identical genetic background, differences in susceptibility reflect compatibility between the phage and pilin receptor. In contrast to the narrow specificity observed for antisera, several phages could infect multiple strains expressing divergent pilins (**Figure 2B, Supplementary Figure S4**). For example, JBD26 and JBD93 infected strains expressing group IIA, IIB, and IIC pilins, despite limited sequence identity among these groups. Notably, DMS3 infected all the pilin library strains except the strain expressing group IV pilins, which are glycosylated mainly on the αβ loop. These findings show that pilin sequence divergence that precludes antibody cross-recognition does not necessarily prevent phage infection, indicating that some phages can tolerate substantial variation in pilin sequence.

### Localized electrostatic perturbations selectively disrupt JBD26 infection

The ability of phages to infect strains expressing divergent pilins suggests that receptor recognition depends on conserved structural or physicochemical features of pilins rather than strict sequence identity. We previously showed that phage DMS3 can infect strains expressing O-antigen glycosylated pilins, whereas the closely related phage JBD26 cannot (28). A chimeric JBD26 phage expressing the distal tail fibres from DMS3 gained the ability to infect glycosylated strains (**Figure 3A**), demonstrating that receptor recognition specificity is determined by those proteins. Therefore, we hypothesized that phages differ in their tolerance to perturbations in pilin surface properties, and phages capable of infecting strains with diverse pilins, such as DMS3, tolerate greater receptor variation.

**Figure 3.**
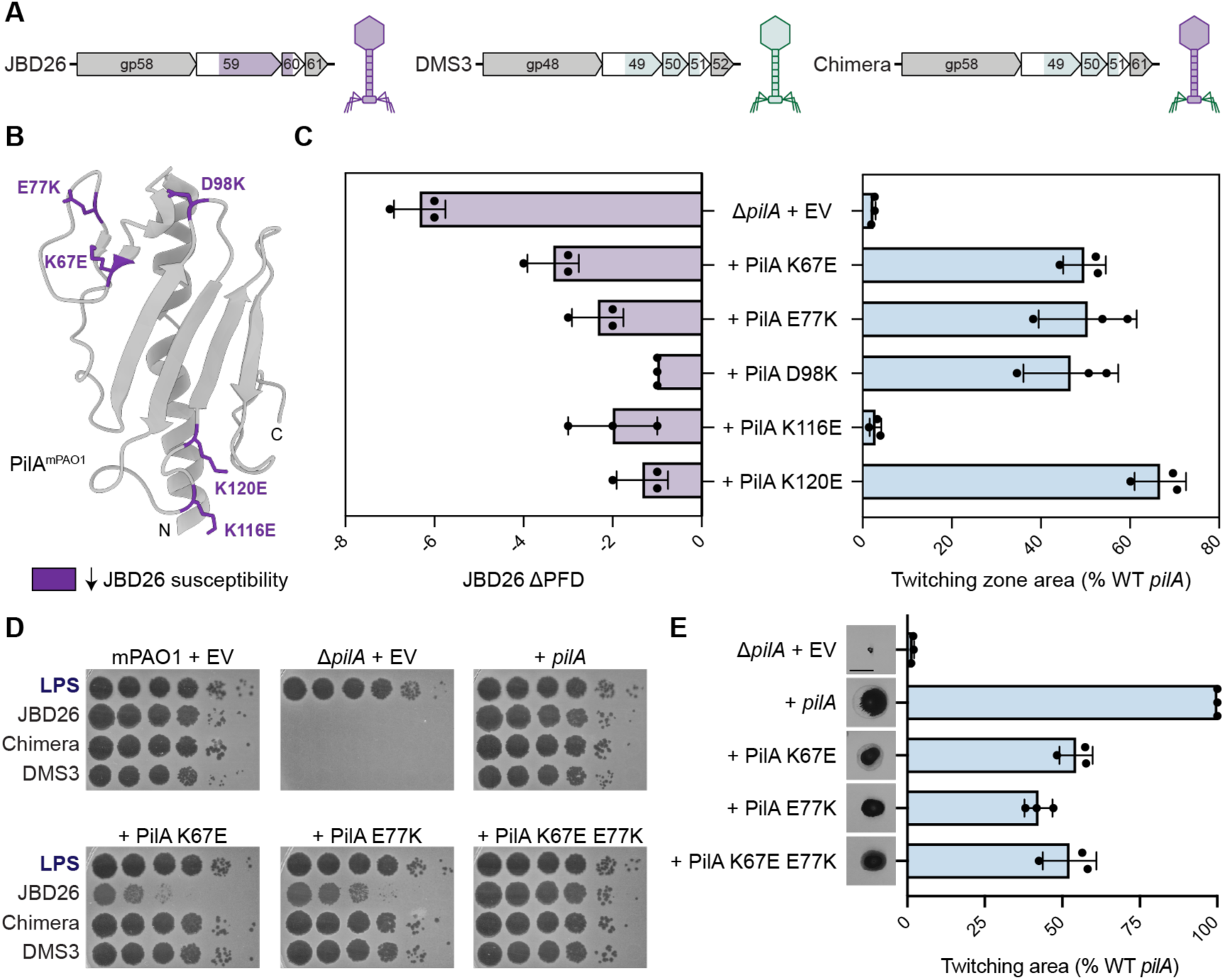
Localized electrostatic perturbations selectively disrupt JBD26 infection. **A.** Organization of tail fibre modules from JBD26, DMS3, and a chimeric JBD26 phage expressing the distal tail fibre and assembly proteins from DMS3. Distal tail fibre gene replacements confer DMS3-like receptor recognition properties (28). **B.** Thirteen charged, solvent-exposed residues in mPAO1 PilA were individually charge-swapped. The five substitutions shown here (K67E, E77K, D98K, K116E, and K120E), mapped onto the mPAO1 PilA structure (PDB 9EWX), reduced JBD26 infectivity. Data for the remaining eight mutants is shown in **Supplementary Figure S5**. **C.** Effects of single charge-swap substitutions on phage infectivity and pilus function. Left, infectivity of JBD26 expressed as the change in plaque-forming dilution (ΛPFD) relative to wild-type *pilA* expressed in the mPAO1 Δ*pilA* background. Right, twitching motility expressed as a percentage of wild-type *pilA* expressed in the mPAO1 Δ*pilA* background. Twitching function was preserved in most mutants. Each point represents an independent experiment. **D.** Plaque assays showing infectivity of phages on strains expressing wild-type *pilA*, single substitutions, or the double mutant (K67E/E77K). Single substitutions reduce infectivity of JBD26 but not DMS3 or the chimera. The double mutant restores JBD26 infectivity. Plaque assays are representative of three independent experiments. **E.** Twitching motility assays showing that pilus function is preserved in single and double mutants, suggesting that altered infectivity reflects receptor recognition rather than loss of pilus assembly. Representative twitching zones and quantification are shown. Twitching assays are representative of three independent experiments. Scale bar, 10 mm.

To test this hypothesis, we introduced charge-swap substitutions in the mPAO1 pilin at solvent-exposed residues predicted to be accessible to phage tail fibres. Charged residues were targeted because electrostatic interactions were previously implicated in RNA phage-pilus binding (14). Thirteen solvent-exposed residues were selected based on the cryo-EM structure of the *P. aeruginosa* mPAO1 pilus filament (38) (**Supplementary Figure S5A**). Mutant strains were tested for susceptibility to JBD26, DMS3, the JBD26 chimera, and an LPS-specific control phage (**Supplementary Figure S5B)**. Twitching motility of each mutant was measured to distinguish defects in phage binding from general impairment of pilus function (**Supplementary Figure S5C**).

Five substitutions – K67E, E77K, D98K, K116E, and K120E – reduced JBD26 infectivity by 10- to 10,000-fold, whereas DMS3 and the JBD26 chimera were minimally affected (0-10-fold change; **Figure 3B, C**). These residues localize to the αβ loop, β1-β2 loop, and β3-β4 loop of PilA (**Figure 3B)**. With the exception of K116E and D129K, most point mutants retained twitching motility (∼30-100% of wild type *pilA*), suggesting that pili remained functional and that reduced infection reflects altered receptor recognition rather than loss of pilus function (**Supplementary Figure S5B)**. Notably, while D129K abolished twitching it did not reduce phage susceptibility, indicating that loss of motility does not necessarily impair infection.

To test whether combining substitutions within the same surface region would further decrease infection, we generated a double mutant (K67E/E77K), as both single substitutions strongly reduced JBD26 susceptibility while retaining pilus function (**Figure 3D**). Surprisingly, although twitching of the double mutant was comparable to the single mutants (∼50% of wild type *pilA*), JBD26 susceptibility was restored to wild-type levels (**Figure 3D, E**). This compensatory effect indicates that receptor recognition by JBD26 depends on the overall electrostatic topology of the pilus surface rather than on individual charged residues. Together, these data show that JBD26 infection is sensitive to perturbations in surface electrostatics within a given structural background, whereas DMS3 tolerates broader variation in pilin surface properties.

### JBD26- and DMS3-like phages encode structurally and evolutionarily distinct tail fibres

The contrasting sensitivities of JBD26 and DMS3 to receptor perturbations indicated that these phages likely engage the pilus using distinct receptor recognition strategies. Because swapping distal tail fibres between these phages transfers tolerance to O-antigen pilin glycosylation and sequence diversification, we hypothesized that differences in tail fibre architecture underlie these phenotypes. To define these differences, we examined the genomic organization, structure, and evolutionary diversity of tail modules from JBD26, DMS3, and related phages.

Comparison of tail module organization revealed a conserved architectural distinction. JBD26 and phages with structurally related tail fibres encode a two-gene tail module consisting of a tail fibre and a single assembly protein (**Figure 4A**). In contrast, DMS3 and phages with structurally related tail fibres encode a three-gene module consisting of a tail fibre followed by two assembly proteins, with one exception having a 2-gene tail module but a DMS3-like tail fibre (**Supplementary File F2**). This difference in gene content suggests that these phage receptor-binding proteins adopt structurally divergent scaffolds.

**Figure 4.**
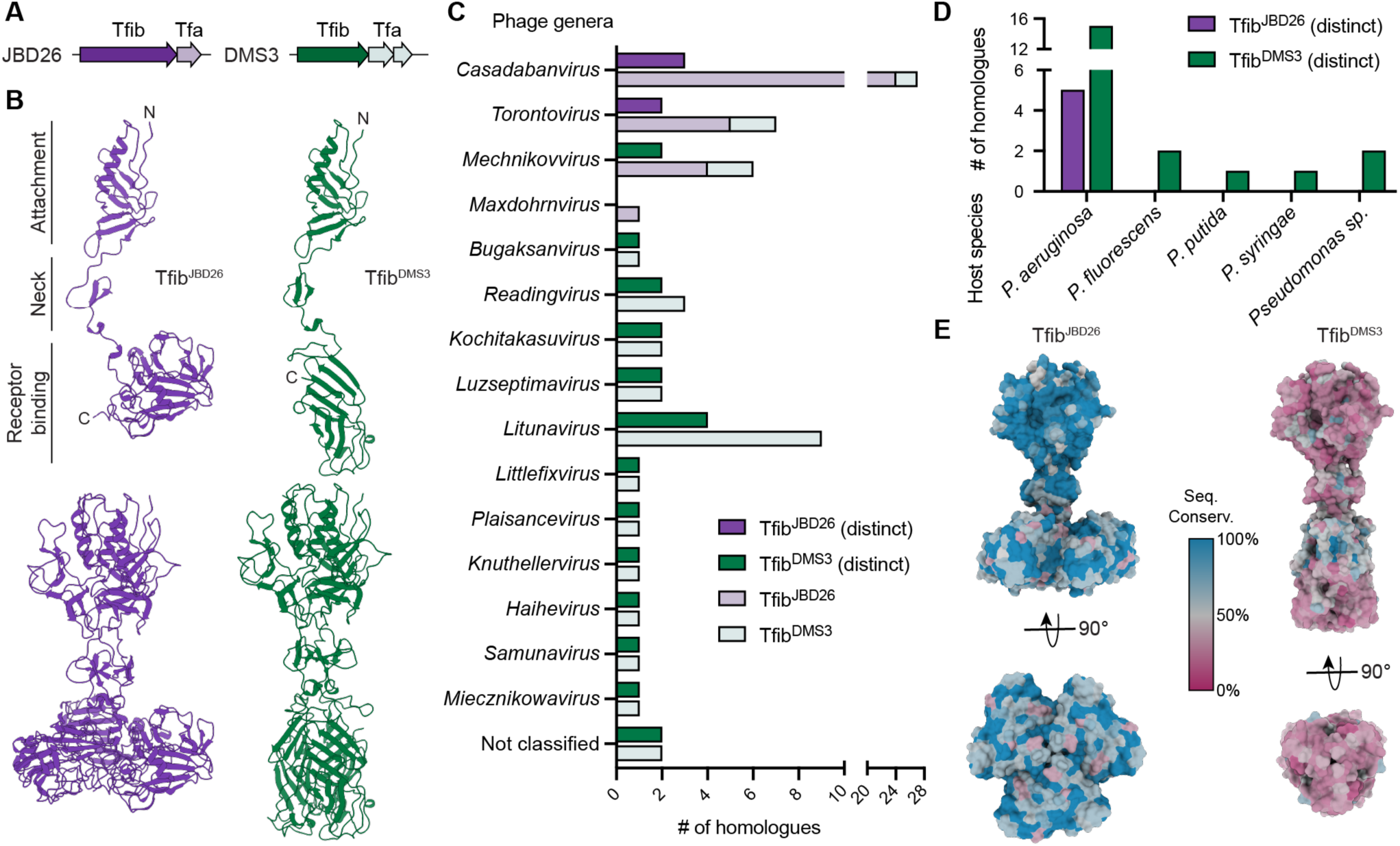
JBD26- and DMS3-like phages encode structurally and evolutionarily distinct tail fibres. **A.** Genomic organization of JBD26- and DMS3-like tail fibre modules. JBD26-like phages encode a tail fibre (Tfib) and single assembly protein (Tfa), whereas DMS3-like phages encode a tail fibre followed by two assembly proteins. **B**. AlphaFold3-predicted (32) structures of JBD26 and DMS3 tail fibres as monomers (top) and in complex (bottom). Both assemble as homotrimers and share a modular organization consisting of an N-terminal attachment domain, neck region, and C-terminal receptor-binding domain. The receptor-binding domains differ substantially in structure, with JBD26-like tail fibres forming compact globular domains and DMS3-like fibres forming elongated β-sandwich domains. AlphaFold3 models coloured by pLDDT values and error plots visualized using UCSF ChimeraX (61) for these structures can be found in **Supplementary Figure S6**. **C.** Distribution of homologous tail fibre modules across phage genera. Tail fibre assembly proteins were used to identify JBD26 and DMS3 tail fibre homologues (JBD26, light purple; DMS3, light green). Tail fibre sequences were then clustered at 90% amino acid sequence identity. DMS3-like tail fibres segregate into more distinct clusters (dark purple) and are distributed across more genera than JBD26-like tail fibres (dark green). **D.** Host species distribution of distinct tail fibre clusters. JBD26-like tail fibres (dark purple) are restricted to phages infecting *P. aeruginosa*, whereas DMS3-like tail fibres (dark green) are found in phages infecting multiple *Pseudomonas* species. **E.** Mapping of sequence conservation onto representative AlphaFold3-predicted structures of tail fibres. JBD26-like fibres (left) exhibit broadly conserved receptor-binding domains, whereas DMS3-like fibres (right) maintain conserved structural cores but diversify extensively at solvent-exposed regions. Residues are coloured according to sequence conservation within tail fibre groups, with conserved residues in blue and variable residues in red.

We next predicted models of these tail fibres using AlphaFold3 (32). Phage tail fibres are typically trimeric assemblies (39), and the homologous receptor-binding protein from phage JBD30 has been structurally resolved as a homotrimer (19). Consistent with this, both JBD26 and DMS3 tail fibres were predicted to assemble as homotrimers, including an N-terminal attachment domain, a connecting “neck” region, and a C-terminal receptor-binding domain (**Figure 4B; Supplementary Figure S6**). Despite this shared modular organization, models of the receptor-binding domains differ substantially. The JBD26 tail fibre terminates in a compact globular domain that forms a broad, disk-like receptor-binding surface, whereas the DMS3 tail fibre adopts a more extended conformation ending in a narrower β-sandwich receptor-binding domain (**Figure 4B**). These structural differences indicate that the two classes of tail fibres present different receptor-interaction surfaces.

To determine whether these tail fibres represent distinct evolutionary lineages, we identified homologous tail modules using the conserved tail fibre assembly proteins as queries against the RefSeq viral protein database (**Supplementary File F2**). Similar numbers of total tail modules were identified for each class (34 for JBD26-like; 32 for DMS3-like), however clustering tail fibre sequences at 90% amino acid sequence identity revealed substantially greater diversification among DMS3-like fibres, which segregated into 21 clusters, compared to only five clusters for JBD26-like fibres (**Figure 4C**, **Supplementary File F2**). Consistent with this difference, DMS3-like tail fibres were identified in phages spanning 13 different genera and associated with multiple *Pseudomonas* species, whereas JBD26-like tail fibres were restricted to two genera and exclusively associated with *P. aeruginosa* (**Figure 4C, D)**.

Mapping sequence conservation onto representative structural models revealed distinct patterns of diversification. JBD26-like fibres showed broad conservation throughout the receptor-binding domain, including exposed surfaces (**Figure 4E**). In contrast, DMS3-like fibres maintained a conserved structural core but exhibited extensive diversification across solvent-exposed regions predicted to contact the receptor. Together, these findings suggest that JBD26-and DMS3-like phages encode structurally and evolutionarily distinct tail fibres.

### Tail module architecture predicts sensitivity to receptor perturbation

If tail fibre architecture dictates receptor recognition strategy, phages encoding similar tail modules should display similar sensitivity to receptor perturbations. To test this hypothesis, we examined additional phages encoding JBD26-like and DMS3-like tail modules. We selected JBD18 and JBD25 (phages with JBD26-like tail modules) and Cootes and Lindberg F10 (phages with DMS3-like tail modules) as representatives. Although these phages encode homologous tail modules, they belong to distinct genera, allowing us to test whether receptor recognition strategy segregates with tail module architecture independent of phage phylogeny.

We first tested whether these phages show similar responses to electrostatic perturbations of the pilus receptor. We exposed K67E, E77K, and the double mutant to each phage (**Figure 5A**). The single mutants displayed reduced susceptibility to JBD26-like phages, with susceptibility restored to near wild-type levels in the double mutant, mirroring the results for JBD26. In contrast, the DMS3-like phages were more tolerant of these substitutions.

**Figure 5.**
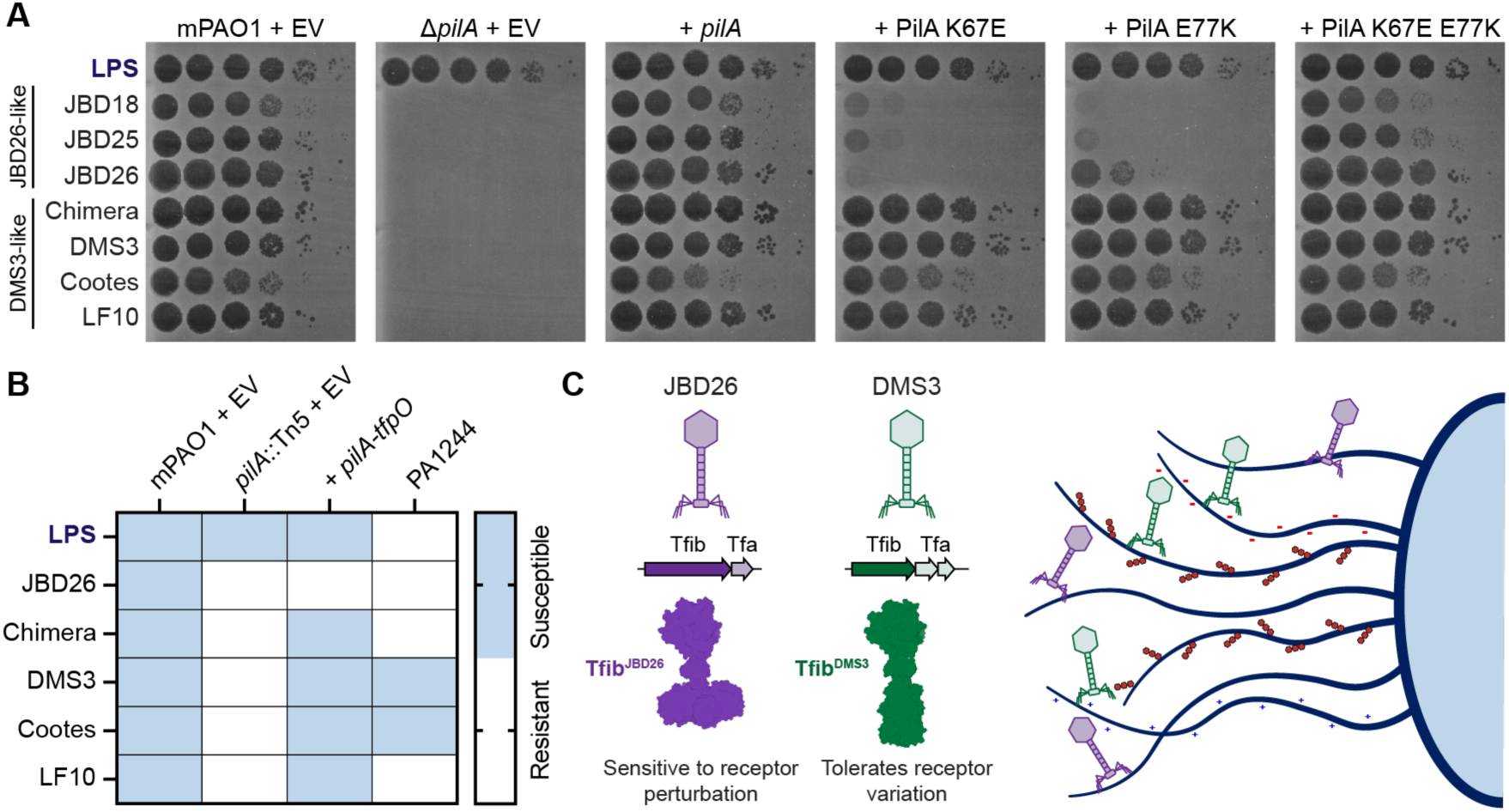
Tail module architecture predicts sensitivity to receptor perturbation. **A.** Plaque assays showing infectivity of representative phages encoding JBD26-like tail modules (JBD18, JBD25, and JBD26) and phages encoding DMS3-like tail modules (JBD26 chimera, DMS3, Cootes, and Lindberg F10 – LF10) on strains expressing wild-type or mutant *pilA* alleles in the mPAO1 Δ*pilA* background. Single substitutions reduce infectivity of JBD26-like phages but not DMS3-like phages. The double mutant restores infectivity of JBD26-like phages. The LPS-targeting phage Kipling (‘LPS’) is shown as a control. Plaque assays are representative of three independent experiments. **B.** Infectivity of phages on strains expressing glycosylated (*pilA-tfpO/*PA1244) or unglycosylated pilins. Glycosylation mediated by TfpO alters pilin surface properties and reduces infectivity of JBD26. All phages with DMS3-like tail fibres retain infectivity in the *pilA-tfpO* recombinant background, whereas only DMS3 and Cootes infect wild-type PA1244. Plaque assays used to generate heat map can be found in **Supplementary Figure S7**. **C.** Model illustrating two different modes of receptor recognition. JBD26-like phages (purple) encode tail fibres with conserved receptor-binding interfaces that require precise electrostatic and structural compatibility with the pilus receptor. In contrast, DMS3-like phages (green) encode structurally distinct tail fibres with diversified receptor-binding surfaces that tolerate receptor variation, enabling infection of pilins with altered sequence or post-translational modification.

We next asked whether this relationship extends to other receptor modifications. Pilin glycosylation introduces bulky post-translational modifications at exposed residues and prevents infection by JBD26 but not DMS3 (28). To test whether tolerance to O-antigen glycosylation also segregates with tail module architecture, we assessed infectivity of JBD26- and DMS3-like phages on strains expressing glycosylated pilins, including a recombinant mPAO1 *pilA* mutant complemented with the *pilA-tfpO* cassette from group I strain PA1244 as well as wild-type strain PA1244 (**Figure 5B**). All DMS3-like phages retained the ability to infect the *pilA-tfpO* recombinant strain, indicating tolerance to glycosylated pilins in this genetic background, whereas JBD26 did not. Infection of wild-type PA1244 was more restricted: only DMS3 and Cootes could infect this strain, likely due to additional strain-specific phage defence factors not present in mPAO1. Together, these results show that sensitivity to electrostatic perturbation and pilin glycosylation segregates with tail module architecture, independently of overall phage phylogeny.

## DISCUSSION

Bacterial surface receptors evolve rapidly, yet phages must adapt to maintain receptor recognition and persist (40). Using the sequence-diverse T4P of *P. aeruginosa* as a model system, we found that pilus-dependent phages differ markedly in their tolerance of receptor diversification. These differences segregate with tail module architecture and are consistent with distinct receptor recognition strategies. Phages encoding structurally conserved receptor-binding tail fibres (JBD26-like) were sensitive to electrostatic perturbations of the pilus surface, whereas phages encoding structurally diversified tail fibres (DMS3-like) tolerated considerable receptor variation (**Figure 5C**). Together, our findings support a model in which tail fibre architecture influences how phages accommodate receptor diversification.

Analysis of more than 1,300 *P. aeruginosa* genomes showed extensive PilA diversification that was concentrated in solvent-exposed regions of the globular domain, while residues required for maintaining the pilin fold and filament assembly remained highly conserved. This pattern permits variation in solvent-accessible surfaces while preserving pilus structure and extension-retraction dynamics (41, 42). Similar strategies occur in other bacteria. For example, *Neisseria* species generate pilin diversity through recombination between expressed and silent pilin loci (43), and *Acinetobacter* species exhibit substantial sequence variation and glycosylation in exposed regions of their pilins (44). These observations indicate that diversification of solvent-exposed pilin surfaces is a common evolutionary strategy that can alter interactions with host and viral binding partners while maintaining pilus function.

Consistent with this concept, natural pilin sequence divergence disrupted polyclonal antibody recognition and prevented infection by a subset of phages. However, several of the phages tested retained infectivity in strains expressing highly divergent pilins. Because all pilins were expressed in a common background, these differences in susceptibility can be attributed to variation in the major pilin or its presentation within the filament. These findings show that some phages can tolerate substantially greater variation than antibodies, consistent with the idea that antibodies recognize specific molecular epitopes, whereas some phages may engage conserved structural or physicochemical features that persist despite local sequence diversification (45, 46).

Targeted mutagenesis revealed that receptor recognition by certain phages is sensitive to localized electrostatic perturbations at the pilus surface. Charge-reversal substitutions within solvent-exposed loops of the globular domain reduced infectivity of JBD26-like phages by several orders of magnitude, while compensatory substitutions restored infection. This result implies that receptor recognition depends on the overall electrostatic topology of the pilus surface rather than strict conservation or arrangement of individual sidechains. Notably, JBD26 could infect strains expressing group IIA, IIB, or IIC pilins despite substantial sequence divergence and local structural variation among these groups. This suggests that compatible physicochemical environments can be achieved within distinct pilin scaffolds, and that electrostatic requirements are satisfied within the three-dimensional context of each variant rather than requiring a fixed electrostatic configuration. Mutations that altered infectivity localized to solvent-exposed loops of the globular domain, particularly the αβ loop, which can tolerate sequence variation and post-translational modification in many pilins (27, 42, 47). This region has also been identified as a determinant of susceptibility for PP7 and related RNA phages (14, 16, 17). In contrast, DMS3-like phages were largely insensitive to these perturbations, indicating that phages differ in their dependence on specific interactions for receptor engagement. Direct adsorption or binding measurements will be required to determine whether altered infectivity reflects changes in initial attachment or downstream steps in the infection process.

Our genetic exchange experiments further support a central role for distal tail fibres in receptor recognition. Swapping distal tail fibre genes between JBD26 and DMS3 altered their sensitivity to pilin charge substitutions and O-antigen glycosylation (28), indicating that these proteins play important roles in determining receptor compatibility. These findings refine prior models proposing roles for proximal tail components in pilus attachment (19), and suggest instead that distal tail fibres are key determinants of receptor specificity and host range.

Structural and evolutionary analyses suggested that these functional differences are associated with intrinsic properties of receptor-binding tail fibres. Although AlphaFold3 (32) models of JBD26- and DMS3-like tail fibres revealed that they share a common modular organization, their terminal receptor-binding domains differ substantially in sequence and structural conservation. While these models have high confidence scores, experimental validation will be required to confirm the details of these architectural differences. Phage tail regions are well-established hotspots of recombination and horizontal exchange, particularly within receptor-binding modules (20–22). Extensive sequence variation within exposed regions could reflect adaptation to receptor change. In such a model, diversification of DMS3-like fibres would reflect evolutionary responses to pilin variation, while conservation of JBD26-like fibres reflects recognition of conserved receptor features.

However, our functional and bioinformatic data do not support this interpretation. Despite their greater sequence diversification, the DMS3-like phages we tested tolerated electrostatic substitutions within group II, large sequence variations between groups, and pilin glycosylation, whereas the more conserved JBD26-like phages were highly sensitive to these perturbations. DMS3-like receptor-binding modules are distributed across more phage genera and associated with infection of multiple *Pseudomonas* species, whereas JBD26-like modules are largely restricted to *P. aeruginosa*. This broader ecological and phylogenetic distribution is consistent with tolerance of receptor variation rather than specialization to narrow receptor sequence space. Thus, diversification of exposed receptor-binding surfaces in DMS3-like modules appears to correlate with expanded receptor compatibility. However, direct structural characterization of tail fibre-pilus complexes will be necessary to define the precise molecular interactions underlying these differences.

Analysis of phages encoding related tail modules showed that tolerance to receptor variation segregates with tail fibre architecture across independent phage lineages. Because these phages represent diverse evolutionary backgrounds, this association suggests that tolerance to receptor variation is linked to tail fibre architecture rather than overall phage relatedness. The distribution across phylogenetically distinct phages may reflect horizontal transfer of tail modules, a common mechanism for reshaping host range in tailed phages, although convergent evolution under similar receptor-selective pressures cannot be excluded (20–22). Tail fibres conferring narrow recognition breadth may constrain host range and limit opportunities for ecological expansion and recombination, whereas architectures allowing broader recognition could facilitate infection of multiple hosts and promote genetic exchange among diverse phages This is consistent with the broader range of host species associated with phages encoding DMS3-like fibres.

While tail module architecture predicts tolerance to receptor variation for phages that engage the major pilin, the behaviour of phages predicted to interact with the minor pilin FimU, such as JBD68 and Lindberg F10 (15), highlights the importance of receptor context within the assembled pilus. FimU is one of five minor pilins that, along with the adhesin PilY1, form a tip complex that primes pilus assembly, and is predicted to link PilA to this complex (48, 49). Both phages have been proposed to engage FimU at the pilus tip; however, all recombinant strains in our pilin library carry identical *fimU* alleles. If FimU sequence alone determined susceptibility, all these strains would be expected to exhibit similar permissivity. Instead, JBD68 infected only strains expressing group II pilins and failed to infect those with O-antigen glycosylated subunits, whereas Lindberg F10 retained infectivity in strains expressing glycosylated pilins. These data indicate that receptor recognition cannot be explained by FimU alone, but instead depends on how receptor components are presented within the filament. Our results suggest that phages proposed to bind FimU may recognize composite structural features formed at the interface between major and minor pilins, reflecting a more complex mode of engagement. Recognition of such composite binding surfaces could help explain the broad host range of phages such as DMS3, although direct imaging by electron microscopy shows that DMS3 particles bind along the length of the pilus filament rather than exclusively at the tip (15).

Together, our findings identify phage tail fibre architecture as a structural correlate of tolerance to variation in T4P receptors. This work provides a structural and evolutionary framework for understanding how phages accommodate receptor evolution. More broadly, these principles can guide both the natural evolution and rational engineering of phages with altered host specificity, important considerations for the development of broad-host-range therapeutic phages to treat antibiotic-resistant infections (49–51).

## METHODS

### Genome collection, MLST assignment, and *pilA* identification

Genome sequences were downloaded from the *Pseudomonas* Genome Database in 2016 (50). To avoid overrepresentation of clonal isolates, closely related genomes from longitudinal or outbreak datasets (including BioProject PRJNA282164 and the Liverpool Epidemic Strain collection) were collapsed to a single representative genome per lineage. Sequence types were assigned using the seven-locus MLST scheme for *P. aeruginosa* (*acsA*, *aroE*, *guaA*, *mutL*, *nuoD*, *ppsA*, *trpE*) as defined by the PubMLST database (51). Alleles were identified using BLASTN searches and assigned according to PubMLST definitions. When novel allele combinations were encountered, phylogenetic context and sequence divergence were used to confirm strain distinctiveness. The final dataset consisted of 1,365 nonredundant genomes representing unique MLST backgrounds.

To identify pilin genes, representative PilA sequences from each of the five established pilin groups were used as queries in BLAST searches against each genome. Candidate *pilA* loci were manually inspected to confirm intact open reading frames and correct genomic context, including proximity to the conserved *pilB* gene. In cases where BLAST searches did not identify a full-length *pilA* gene, the conserved N-terminal coding region and surrounding genomic loci were examined manually to identify divergent pilin sequences. Pilin sequences containing frameshifts or premature stop codons were excluded. Mature pilin sequences were aligned using MUSCLE (52), and identical sequences present in multiple isolates were collapsed to generate a nonredundant set of pilin alleles. Pilin alleles were assigned to groups based on accessory gene content and C-terminal disulfide-bonded loop length (24). A total of 53 distinct mature PilA sequences representing 319 unique pilin-MLST combinations were identified (**Supplementary File F1**).

### Phylogenetic analysis of group II pilins

Unique group II pilin sequences were aligned using Clustal Omega (1.2.3) (53). Maximum likelihood phylogenetic inference was performed using IQ-TREE 2 (2.2.6) (54) with automatic model selection (‘-m MFP’). Branch support was assessed using ultrafast bootstrap approximation with 1,000 replicates (‘-B 1000’). Trees were visualized and annotated using iTOL (7.4.2) (55).

### Bacterial strains, plasmids, and phages

Bacterial strains, plasmids, and phages used in this study are listed in **Supplementary Table S3**. *E. coli* and *P. aeruginosa* strains were grown in lysogeny broth (LB; BioShop) at 37°C with shaking at 200 rpm or on 1.5% LB agar supplemented with antibiotics as required. Antibiotics were used at the following concentrations (µg/mL): ampicillin, 100; carbenicillin, 200; gentamicin, 15 for *E. coli* and 30 for *P. aeruginosa*. Arabinose-inducible constructs were expressed using 0.02-0.2% L-arabinose.

Plasmids were introduced into chemically competent *E. coli* by heat shock and into *P. aeruginosa* by electroporation. Constructs were verified by Sanger sequencing (McMaster Genomics Facility) or by Oxford Nanopore Technology whole-plasmid sequencing (Plasmidsauras).

### Structural determination of PilA^1244^

His_6_-V5-PilA^1244^ lacking the N-terminal signal peptide (Δ1-28, mature protein) was cloned into the expression vector pET151 and transformed into *E. coli* Origami (DE3). Protein expression was induced with 0.5 mM IPTG at 16°C overnight. Cells were harvested by centrifugation (3,200 x g, 4°C, 15 min) and resuspended in lysis buffer (20 mM Tris-HCl, pH 8.0, 500 mM NaCl, 0.1% lauryl dimethylamine N-oxide). Cells were lysed by sonication, and the lysate was clarified by centrifugation (48,258 x g, 4°C, 45 min).

The supernatant was applied to a 5-ml Ni Sepharose High Performance column (GE Healthcare) pre-equilibrated with Ni buffer A (20 mM Tris-HCl, pH 8.0, 500 mM KCl, 10 mM imidazole) using an AKTA Start fast protein liquid chromatography (FPLC) system. The column was washed stepwise with buffer containing 25, 40, and 55 mM imidazole, and bound protein was eluted with Ni buffer B (20 mM Tris-HCl, pH 8.0, 500 mM KCl, 300 mM imidazole).

Eluted fractions were dialyzed overnight at 4°C in 20 mM Tris-HCl, pH 8.0, 150 mM NaCl in the presence of TEV protease (1 mg) to remove the His_6_ and V5 epitope tags. The cleaved sample was reapplied to the Ni Sepharose column, and the untagged PilA^1244^ protein was collected in the flow-through.

Purified PilA^1244^ was concentrated and dialyzed into 20 mM Tris-HCl, pH 8.0, 50 mM NaCl. Crystallization trials were performed using the MCSG Suite (Anatrace). Hanging drops consisting of 1 µl protein solution mixed with 1 µl reservoir solution were dehydrated over 500 µl of 1.5 M ammonium sulfate at 22°C. Crystals formed after approximately four months under conditions containing 0.1 M HEPES-NaOH (pH 7.5), 20% (w/v) PEG 4000, and 10% (v/v) 2-propanol. PilA^1244^ crystals were cryo-protected by addition of 20% ethylene glycol and flash frozen in liquid nitrogen.

All X-ray diffraction data sets were collected using a home source generator (Rigaku MicroMax-007 HF equipped with RAXIS-IV++ detector) and subsequently indexed, processed, and merged using HKL3000 (56). An initial molecular replacement (MR) solution was obtained using the group II pilin from *P. aeruginosa* strain PAK (PDB ID: 1OQW, residues 29-144) (57) as a search template, modified to poly-alanine model using PDB file editor in the PHENIX suite (58). However, model building and refinement was hindered by model bias. To improve the initial phases, we exploited the presence of two highly conserved C-terminal Cys residues flanking the disulfide-bonded loop. We performed MR-SAD (59) using the partial MR solution to find two sulfur sites with figure of merit (FOM) = 0.438 and occupancies of 0.75 and 0.84, respectively. A complete model was built using AutoBuild with the improved phase information, which was improved through iterative cycles of manual model building in COOT (60) and refinement in PHENIX (**Supplementary Table S2**).

### AlphaFold3 modeling and structural analysis

Protein structures were predicted using AlphaFold3 (32). Model confidence was assessed using predicted local distance difference test (pLDDT) scores and predicted aligned error (PAE) plots. Structures were visualized and analyzed using UCSF ChimeraX (1.7.1) (61). Sequence alignments were generated using Clustal Omega (1.2.3) (53). Conservation scores were calculated based on amino acid sequence identity and mapped onto structural models using UCSF ChimeraX (1.7.1). Solvent accessibility for PilA mutagenesis experiment was inferred based on structural visualization and comparison with the cryo-EM structure of the mPAO1 pilus (PDB 9EWX).

### Construction of pilin expression library

Representative *pilA* genes from group I (synthesized by IDT) and groups IIB and IIC (synthesized by GenScript) were cloned into the arabinose-inducible vector pBADGr using EcoRI and HindIII restriction digestion followed by ligation and transformation into *E. coli* DH5α. Constructs were transformed into a mPAO1 *pilA*::Tn5 mutant. PilA expression for pilin library experiments was induced with 0.02% L-arabinose.

### Twitching motility assays

Twitching motility was tested as previously described (62). In brief, overnight single colonies were stab-inoculated in triplicate to the bottom of cell culture-treated Nunc™ OmniTray™ Single-Well Plates (ThermoFisher Scientific) containing 1% LB agar, supplemented with L-arabinose and antibiotic when necessary. After inoculation, plates were incubated at 37°C for 16-18 h. Agar was carefully removed after incubation and twitching zones were stained with 1% crystal violet for 10 min. Plates were rinsed with distilled water to remove excess dye and air-dried. Plates were imaged using a flatbed scanner and twitching zone area was measured using ImageJ (63).

### Sheared surface protein preparation and SDS-PAGE analysis

Sheared surface proteins were collected as previously described with modifications (34). Briefly, strains of interest were streaked in a grid pattern on 1.5% LB agar plates supplemented with gentamicin and L-arabinose and incubated at 37°C for ∼16 h. Cells were scraped from the plates with glass coverslips and resuspended in 4.5 mL of 1X sterile phosphate-buffered saline (PBS; pH 7.4). Surface proteins were sheared by vortexing the cell suspensions for 30 s. The suspensions were divided into three 1.5 mL microcentrifuge tubes, and cells were pelleted by centrifugation at 21,000 x g for 1 h. Supernatants were transferred to fresh tubes, and surface proteins were precipitated by adding 1/10 volume of 5 M NaCl and 30% (w/v) polyethylene glycol (PEG 8000) to each tube and incubating on ice for 90 min. Precipitated proteins were collected by centrifugation at 21,000 x g for 30 min and resuspended in 150 µL 1X sodium dodecyl sulfate polyacrylamide gel electrophoresis (SDS-PAGE) sample loading dye (125 mM Tris-HCl, pH 6.8; 2% β-mercaptoethanol; 20% glycerol; 4% SDS; 0.001% bromophenol blue). Samples were pooled, boiled for 10 min, centrifuged at 21,000 x g for 10 min, and resolved on 15% SDS-PAGE gels at 120 V in 1X Tris-glycine running buffer with a prestained protein ladder (BLUelf). Protein bands were visualized using Coomassie blue staining.

### Antibody cross-reactivity assays

After vortexing cells to shear surface pili and flagella, the remaining cells were resuspended in sterile 1X PBS to a final optical density (OD_600_) of 0.6. One mL of this standardized culture was transferred to a microcentrifuge tube and cells were pelleted at 21,000 x g for 10 min. The supernatant was removed, and the pelleted cells were resuspended in 50 μL of 1X SDS-PAGE sample loading dye. Samples were boiled for 10 min, centrifuged for 10 min at 21,000 x g, and then resolved on 15% SDS-PAGE gels at 120 V in 1X Tris-glycine running buffer with a prestained protein ladder (BLUelf). After SDS-PAGE, proteins were transferred to a 0.45 μm nitrocellulose membrane (BioRad) for 1 h at 225 mA in 1X transfer buffer (20% methanol and 100 mL Tris-glycine buffer stock without SDS in 1 L Milli-Q H_2_O). Membranes were blocked with 5% skim milk (BioRad) resuspended in 1X PBS for 2 h at room temperature. For detection of PilA, blots were incubated with 1/5000 dilution (1X PBS) of rabbit polyclonal antibodies, as described previously (24). Antisera for groups I (this study; Cedarlane), II (33), and IV (27) were generated against N-terminally truncated recombinant pilins expressed in *E. coli*, whereas antisera for groups III and V (34) were raised against sheared surface protein preparations from *P. aeruginosa*. Blots were incubated with primary antibodies overnight at room temperature. Membranes were washed with 1X PBS and then incubated with 1/3000 dilution (1X PBS) of goat α-rabbit alkaline phosphatase conjugated secondary antibodies (BioRad) for 1 h at room temperature. Membranes were washed with 1X PBS and then developed using 5-bromo-4-chloro-3-indolylphosphatase (BioShop) and nitro blue tetrazolium (BioShop) resuspended in alkaline phosphatase buffer (1 mM Tris, 100 mM NaCl, 5 mM MgCl_2_, pH 9.5).

### Phage propagation and titration

Phages were propagated on *P. aeruginosa* mPAO1. Overnight cultures were subcultured 1:100 into LB medium and grown to an OD_600_ of 0.3-0.5. Cultures were then infected with phage at an MOI of 0.1 and incubated at 37°C with shaking at 200 rpm until complete lysis was observed. Lysates were clarified by centrifugation at 21,000 x g for 10 min to remove cellular debris and subsequently filtered through sterile 0.45 µm filters to remove residual bacterial cells. Phage titres were determined by plaque assay using serial dilution and expressed as plaque-forming units per milliliter (PFU/mL).

### Phage isolation and genome sequence analysis

Phage isolation and genome sequence analysis were performed as described previously for phages Kipling and Cootes (64). BLASTN analysis showed that phages Kipling and Cootes shared ∼97% sequence identity with *Pseudomonas* phages vB_PaeM_CEB_DP1 (NC_041870) and KPP25 (NC_024123), respectively.

### Phage plaquing assays

Phage plaque assays were performed as previously described (28), with modifications. Briefly, bacteria were grown overnight at 37°C in LB supplemented with gentamicin with shaking at 200 rpm, then subcultured 1:100 into LB containing 10 mM MgSO_4_, gentamicin, and L-arabinose and incubated at 37°C with shaking for 3 h. Cultures were standardized to an OD_600_ of 0.3 in LB containing 10 mM MgSO_4_, and 100 µL was mixed with 12 mL molten top agar (LB, 10 mM MgSO_4_, 0.6% agar) and overlaid onto pre-poured rectangular (Nunc™ OmniTray™ Single-Well Plates) LB agar plates (1.5% agar, 10 mM MgSO_4_) supplemented with antibiotics and L-arabinose where indicated. After solidification, plates were air-dried in a biosafety cabinet for 30 min.

Phage stocks standardized to 10^8^ PFU/mL were serially diluted 10-fold in LB containing 10 mM MgSO_4_, and 5 µL of each dilution was spotted onto the prepared plates. After air-drying for 10 min, plates were incubated at 37 °C for 18 h. Plates were imaged, and phage titre was estimated as the lowest dilution yielding countable plaques, referred to as the plaque-forming dilution (PFD). Reported PFD values represent the mean of three independent biological replicates.

### Site-directed mutagenesis of *pilA*

Site-directed mutagenesis was performed using overlap extension PCR with the pHERD30T-*pilA* plasmid as the template to introduce point mutations into the pilin coding sequence (65). All primers used in this study are listed in **Supplementary Table S4**. Mutant *pilA* fragments were cloned into the arabinose-inducible vector pHERD30T using EcoRI and HindIII restriction digestion followed by ligation and transformation into *E. coli* DH5α. Constructs were transformed into a mPAO1 Δ*pilA* mutant. Expression of *pilA* variants was induced with 0.2% L-arabinose.

### Comparative analysis of phage tail modules

Tail fibre assembly proteins from *Pseudomonas* phages DMS3 (gp50 and gp51; YP_950474.1, YP_950475.1) and JBD26 (gp60; YP_010299254.1) were used as queries in BLASTP searches against the NCBI RefSeq viral protein database (downloaded February, 2026) using BLAST+ (2.13.0) with an E-value threshold of 1x10^-5^. Hits were retained if they aligned over ≥50% of the query length with ≥30% amino acid identity. Corresponding genome accessions were retrieved using Entrez Direct (24.0), and GenBank files were downloaded and parsed using Biopython (1.86). Genomic regions spanning 10 coding sequences upstream and downstream of each assembly protein gene were extracted.

Tail fibre genes and module architecture were identified by manual inspection in Geneious Prime (2024.0) based on gene length, annotation, and genomic context. Tail fibre sequences were clustered using CD-HIT (4.8.1) (66) at 90% amino acid sequence identity to define distinct tail fibres. Phage taxonomy and host annotations were obtained from GenBank metadata, and the most frequent annotation was assigned to each cluster.

Diversity was quantified as the number of tail fibre clusters, while taxonomic breadth, host breadth, and ecological prevalence were defined as the number of associated phage genera, host species, and total tail modules, respectively. Representative structures for each cluster were predicted using AlphaFold3 (32) and classified based on structural similarity to JBD26-like and DMS3-like tail fibres. All database accessions and identifiers are provided in **Supplementary File F2**.

### Data visualization

Graphs were generated using GraphPad Prism (10.5.0). Structures were visualized using UCSF ChimeraX (1.7.1) (61).

### Data availability

Structural data have been deposited in the Protein Data Bank under accession number 6BBK. All sequence identifiers and analysis outputs are provided in **Supplementary File F1** and **Supplementary File F2**. Annotated phage genome sequences were submitted to GenBank under accession numbers PZ053911 (*Pseudomonas* phage Kipling) and PZ053910 (*Pseudomonas* phage Cootes).

## ACKNOWLEDGEMENTS

We thank Yi Ni Liu for assistance with generating the initial pilin-MLST dataset; Dr. Karen Maxwell for providing select *Pseudomonas* phages; Dr. Alexander Hynes for access to the Hynes Lab Bioinformatics Workstation; and Dr. Alan Davidson and members of the Burrows Lab for helpful discussions. Biren Dave, Brandon Aubie, Rachel Tran, and Anna-Lisa Nguyen performed exploratory analyses leading to final conceptualization of this work. This work was funded by Canadian Institutes of Health Research Project grants (PJT-156080) to LLB, (PJT-189946) to AG, and (PJT-156214) to AGM. LLB holds a Tier 1 Canada Research Chair in Microbe-Surface Interactions (CRC 2021-00103), AG holds a Tier 1 Canada Research Chair in DNA Transposition (CRC 2024-00369), and AGM held a Cisco Research Chair in Bioinformatics and holds a David Braley Chair in Computational Biology. IQ holds a doctoral Natural Sciences and Engineering Research Council Canada Graduate Scholarship and IC was supported by a David Braley Centre for Antibiotic Discovery Summer Studentship.

