## Supplementary Figures S1-S7 and Tables S1-S4 for "Structural diversification of phage tail fibres enables recognition of diverse type IV pili"

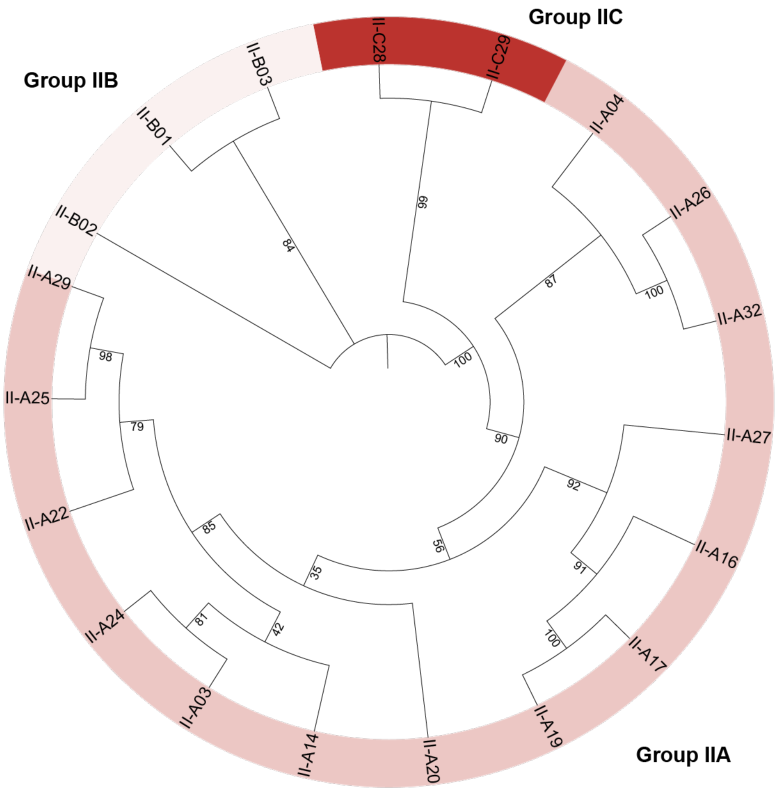


**Supplementary Figure S1. Phylogenetic analysis of distinct subgroups within group II pilins.** Unique group II pilin sequences were aligned using ClustalW (1), phylogenetic relationships were inferred with IQ-TREE 2 (parameters used '-m MFP -B 1000') (2), and the resulting tree was visualized using iTOL (3). Bootstrap values are indicated at each branch. The tree resolves group II pilins into three distinct subgroups, corresponding to the previously defined group IIA and the newly identified subgroups IIB and IIC (coloured in three shades of red).

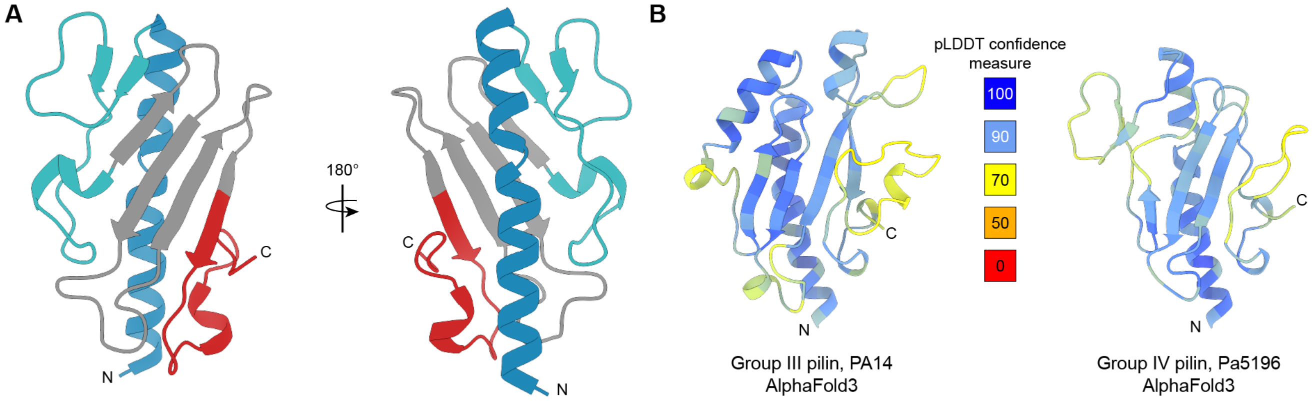


**Supplementary Figure S2. Structural context and model confidence for representative pilins. A.** Crystal structure of the N-terminally truncated group I pilin $\text{PilA}^{1244}$, shown in two orientations related by a 180° rotation. The structure adopts the canonical type IV pilin fold, consisting of an extended N-terminal α-helix (blue), an αβ loop (cyan), a four-stranded antiparallel β-sheet (grey), and a C-terminal disulfide-bonded D-region (red). The structure was determined at 1.7 Å resolution using a construct lacking the N-terminal membrane-associated segment (Δ1-28). Data collection and refinement statistics are provided in **Supplementary Table S2**. **B.** AlphaFold3 (4) models of representative group III (PA14) and group IV (Pa5196) pilins coloured by predicted local distance difference test (pLDDT) confidence scores.

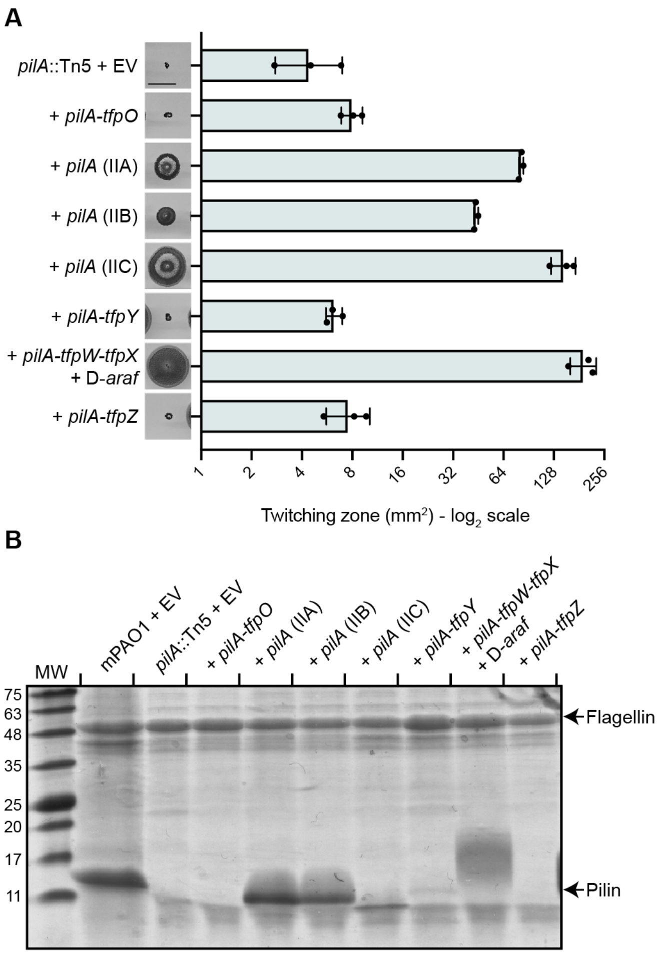


**Supplementary Figure S3. Pilin expression and function in recombinant strains. A.** Twitching motility assays of strains expressing representative pilins in the mPAO1 *pilA* mutant background. Pilin alleles were expressed from plasmids together with their associated accessory proteins required for pilin assembly or modification. For group IV pilins, genes required for D-arabinofuranose (D-*araf*) biosynthesis and transfer were coexpressed to enable pilin glycosylation. Twitching zone area was quantified and plotted on a $log_2$ scale. Images are representative of three independent experiments. Scale bar, 10 mm. **B.** SDS-PAGE analysis of sheared surface pili preparations from the same strains. Coomassie-stained gels show PilA (~15 kDa) and flagellin (~50 kDa), which was used as a loading control. Gel is representative of two independent experiments.

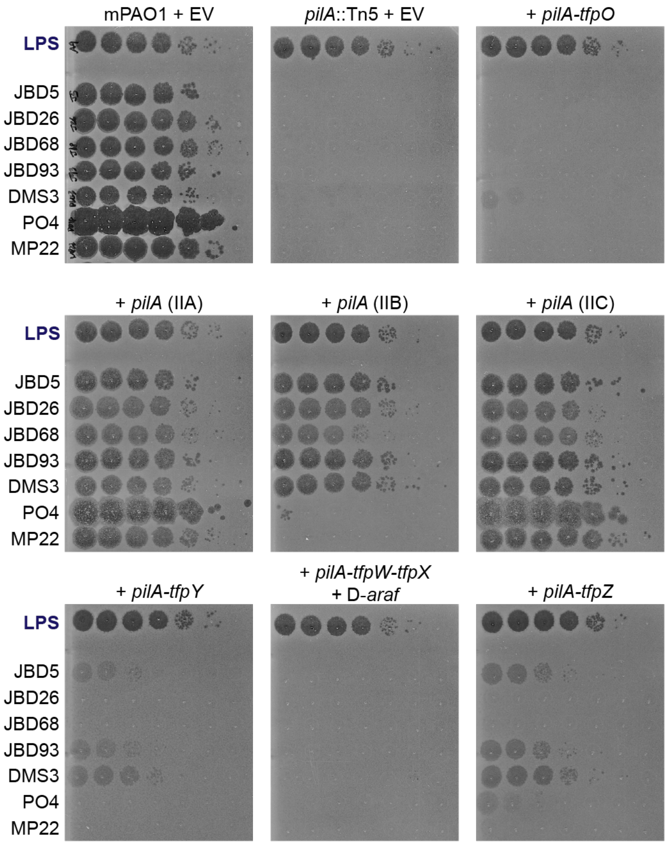


**Supplementary Figure S4. Pilus-dependent phages show differential infectivity across strains expressing divergent pilins.** Representative plaque assays of pilus-dependent phages (JBD5, JBD26, JBD68, JBD93, DMS3, PO4, and MP22) and the lipopolysaccharide (LPS)-targeting control phage Kipling ('LPS') on mPAO1 *pilA* mutant strains complemented with the indicated *pilA* alleles. Pilins were expressed from plasmids together with their associated accessory proteins required for pilin assembly or modification. For group IV pilins, genes required for D-arabinofuranose (D-*araf*) biosynthesis and transfer were coexpressed to enable pilin glycosylation. Phages were serially diluted 10-fold and spotted onto bacterial lawns. Plaque assays are representative of three independent experiments and were used to calculate plaque-forming dilution (PFD) values shown in **Figure 2B**.

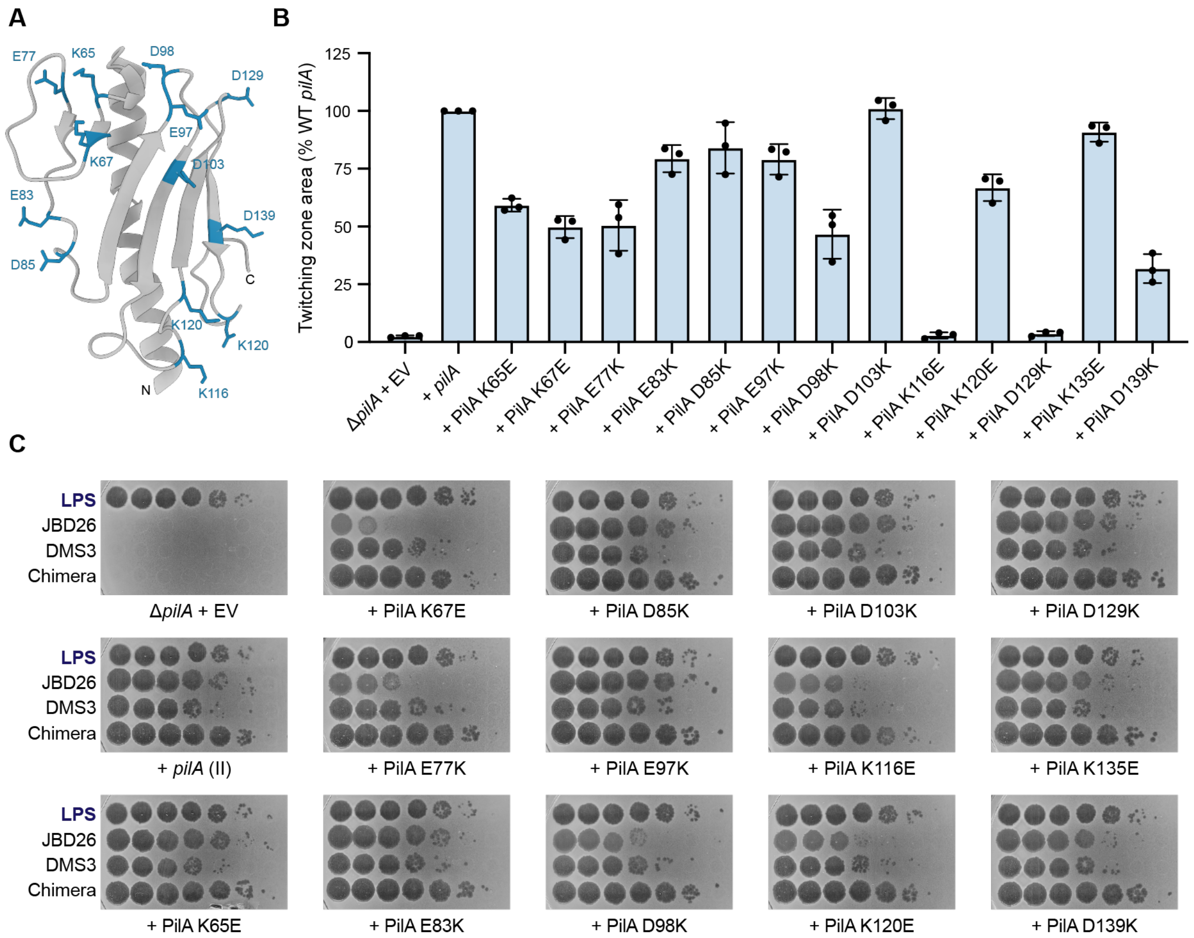


**Supplementary Figure S5. Functional analysis of charge-swap substitutions at solvent-exposed residues of PilA. A.** Thirteen solvent-exposed residues selected for charge-swap mutagenesis mapped onto the mPAO1 PilA cryo-EM structure (PDB 9EWX). **B.** Twitching motility of strains expressing PilA charge-swap mutants in the mPAO1 Δ*pilA* background, expressed as a percentage of wild-type *pilA*. Most substitutions retain twitching motility, indicating that pili remain functional. Each point represents an independent experiment. **C.** Infectivity profiles of JBD26, DMS3, the JBD26 chimera, and the LPS-specific control phage Kipling ('LPS') across all mutants. DMS3 and the chimera retain infectivity across most mutants, whereas JBD26 shows selective sensitivity to specific PilA substitutions. Plaque assays are representative of two independent experiments.

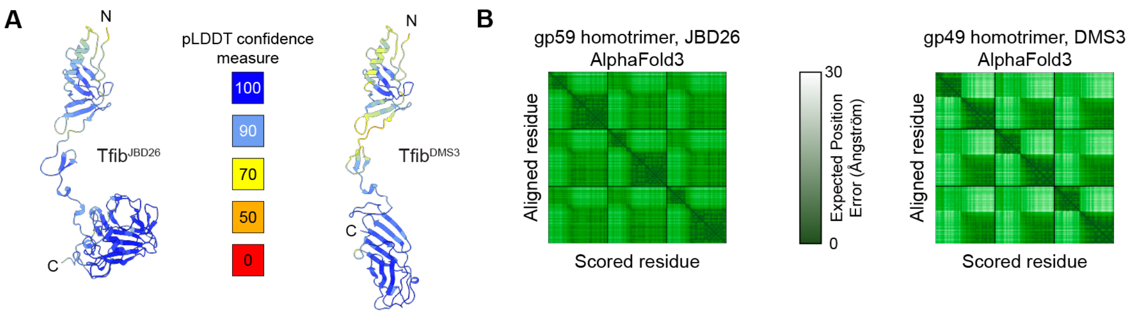


**Supplementary Figure S6. AlphaFold3 model confidence of JBD26-like and DMS3-like tail fibres. A.** AlphaFold3 (4) models of JBD26 tail fibre gp59 (Tfib$^{JBD26}$) and DMS3 tail fibre gp49 (Tfib$^{DMS3}$) coloured by pLDDT confidence scores. **B.** Predicted aligned error (PAE) plots for homotrimers of tail fibre proteins from JBD26 (gp59) and DMS3 (gp49), generated using AlphaFold3.

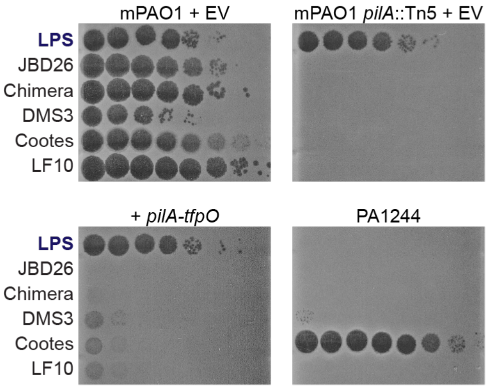


**Supplementary Figure S7. Plaque assays showing infectivity of phages on strains expressing glycosylated or non-glycosylated pilins.** Representative plaque assays demonstrating infectivity of pilus-dependent phages on strains expressing pilins with or without glycosylation. Strains include the glycosylated PA1244 background and the mPAO1 *pilA* mutant complemented with *pilA* and the glycosyltransferase *tfpO*. Phages tested include the LPS-targeting control phage Kipling ('LPS'), JBD26, the JBD26 chimera, and DMS3-like phages Cootes and Lindberg F10 (LF10). These data correspond to the infectivity summary shown in **Figure 5B**. Plaque assays are representative of three independent experiments.

**SUPPLEMENTARY TABLES**

**Supplementary Table S1. Amino acid sequence similarity within and between PilA groups.**

| **Group** | I | II (all)[a] | IIA | IIB | IIC | III | IV | V |
|---|---|---|---|---|---|---|---|---|
| I | 81.2[b] | | | | | | | |
| II (all) | 36.2 | 58.9 | | | | | | |
| IIA | 36.3 | N/A | 77.5 | | | | | |
| IIB | 32.6 | N/A | 32.7 | 98.7 | | | | |
| IIC | 40.8 | N/A | 38.1 | 32.1 | 98.6 | | | |
| III | 27.6 | 24.7 | 24 | 27.1 | 25.6 | 97.5 | | |
| IV | 32.1 | 30.3 | 28.9 | 36.9 | 30.6 | 29.8 | 86.9 | |
| V | 26.5 | 24.8 | 24.7 | 26.6 | 22.7 | 49.9 | 27.3 | 99.2 |

[a] Average similarity among all group II pilins including IIA, IIB, IIC

[b] Average similarity colour-coded from highest (darkest) to lowest (lightest)

**Supplementary Table S2. X-ray data collection and refinement statistics.**

| Crystal | N-terminally truncated PilA$^{1244}$ |
|---|---|
| ***Data collection*** | |
| **Space group** | $P2_12_12_1$ |
| **Unit cell parameters (Å)** | $a$ = 30.5<br>$b$ = 42.4<br>$c$ = 82.4 |
| **X-ray source** | Rotating anode |
| **Wavelength (Å)** | 1.5418 |
| **Resolution (Å)** [a] | 35.00-1.70 (1.73-1.70) |
| **Unique reflections** | 12169 (801) |
| **Average redundancy** | 35.6 (11.0) |
| **Completeness (%)**[a] | 98.8 (78.2) |
| $I/\sigma(I)$ [a] | 88.1 (6.1) |
| $R_{meas}$ **(all) (%)**[a] | 0.052 (0.228) |
| $R_{p.i.m.}$ **(all) (%)**[a] | 0.008 (0.060) |
| ***Refinement*** | |
| **Resolution (Å)** [a] | 29.55-1.73 |
| $R_{work}$ **(%)** | 18.3 |
| $R_{free}$ **(%)** | 21.7 |
| **No. of protein atoms refined** | 962 |
| **No. of solvent atoms refined** | 122 |
| **Mean B factors ($Å^2$)** | 21.2 |
| **Ramachandran statistics (%)** | |
| **Favored / Allowed / Outliers** | 99.2 / 0.8 / 0 |
| **RMSD bond lengths (Å)** | 0.005 |
| **RMSD bond angles (°)** | 0.747 |
| **PDB accession code** | 6BBK |

(a) Values in parentheses correspond to the highest resolution shell

**Supplementary Table S3. Plasmids, bacterial strains, and bacteriophages used in this study.**

| Plasmid/strain/phage | Characteristics | Source |
|---|---|---|
| **Plasmids** | | |
| pET151-1244 *pilA* | Expression vector with N-terminally truncated (Δ 1-28 mature pilin) PA1244 PilA | This work |
| pBADGr | Arabinose-inducible complementation vector | (5) |
| pBADGr-*pilA-tfpO* | pBADGr expressing PA1244 *pilA-tfpO* | (5) |
| pBADGr-*pilA* (IIA) | pBADGr expressing mPAO1 *pilA* | (5) |
| pBADGr-*pilA* (IIB) | pBADGr expressing IIB-03 *pilA* | This work |
| pBADGr-*pilA* (IIC) | pBADGr expressing IIC-29 *pilA* | This work |
| pBADGr-*pilA-tfpY* | pBADGr expressing PA14 *pilA-tfpY* | (5) |
| pBADGr-*pilA-tfpW-tfpX* | pBADGr expressing Pa5196 *pilA-tfpW-tfpX* cassette | (5) |
| pBADGr-*pilA-tfpZ* | pBADGr expressing Pa1457 *pilA-tfpZ* cassette | (5) |
| pBADGr-*pilA* (PA1244) | pBADGr expressing PA1244 *pilA* (group I pilin) | (6) |
| pBADGr-*pilA* (PA13756) | pBADGr expressing AZPAE13756 *pilA* (group I pilin) | This work |
| pBADGr-*pilA* (PA13877) | pBADGr expressing AZPAE13877 *pilA* (group I pilin) | This work |
| pBADGr-*pilA* (PA14868) | pBADGr expressing AZPAE14868 *pilA* (group I pilin) | This work |
| pUCP20 | Broad-host-range cloning vector | (7) |
| pUCP20-D-*araf* | pUCP20 expressing the PA5196 PsPA7_6245-51 genes | (8) |
| pHERD30T | Arabinose-inducible complementation vector | (9) |
| pHERD30T-*pilA* | pHERD30T expressing mPAO1 *pilA* | This work |
| pHERD30T-PilA K65E | pHERD30T expressing mPAO1 PilA K65E | This work |
| pHERD30T-PilA K67E | pHERD30T expressing mPAO1 PilA K67E | This work |
| pHERD30T-PilA E77K | pHERD30T expressing mPAO1 PilA E77K | This work |
| pHERD30T-PilA E83K | pHERD30T expressing mPAO1 PilA E83K | This work |

| | | |
|---|---|---|
| pHERD30T-PilA D85K | pHERD30T expressing mPAO1 PilA D85K | This work |
| pHERD30T-PilA E97K | pHERD30T expressing mPAO1 PilA E97K | This work |
| pHERD30T-PilA D98K | pHERD30T expressing mPAO1 PilA D98K | This work |
| pHERD30T-PilA D103K | pHERD30T expressing mPAO1 PilA D103K | This work |
| pHERD30T-PilA K116E | pHERD30T expressing mPAO1 PilA K116E | This work |
| pHERD30T-PilA K120E | pHERD30T expressing mPAO1 PilA K120E | This work |
| pHERD30T-PilA D129K | pHERD30T expressing mPAO1 PilA D129K | This work |
| pHERD30T-PilA K135E | pHERD30T expressing mPAO1 PilA K135E | This work |
| pHERD30T-PilA D139K | pHERD30T expressing mPAO1 PilA D139K | This work |
| pHERD30T-PilA K67E E77K | pHERD30T expressing mPAO1 PilA K67E E77K. PilA K67E used as template. | This work |
| ***E. coli* strains** | | |
| DH5α | *F⁻φ80lacZΔM15 Δ(lacZYA-argF)U169 recA1 endA1 hsdR17(rk⁻, mk⁺) phoA supE44 thi-1 gyrA96 relA1 λ−* | Invitrogen |
| Origami (DE3) | F-*ompT hsd*SB(rB- mB-) *gal dcm lacY1 ahpC* (DE3) *gor522*::Tn10 *trxB* ($Kan^R$, $Te^{tR}$) | Novagen |
| ***P. aeruginosa* strains** | | |
| mPAO1 | WT | (10) |
| mPAO1 + EV | WT complemented with pBADGr/pHERD30T | (5) |
| mPAO1 *pilA*::Tn5 | Tn5-phoA insertion at base 163 in *pilA* | (10) |
| mPAO1 *pilA*::Tn5 + EV | *pilA* transposon mutant complemented with pBADGr | (5) |
| mPAO1 *pilA*::Tn5 + *pilA-tfpO* | *pilA* transposon mutant complemented with pBADGr-*pilA-tfpO* | (5) |
| mPAO1 *pilA*::Tn5 + *pilA* (IIA) | *pilA* transposon mutant complemented with pBADGr-*pilA* (IIA) | |
| mPAO1 *pilA*::Tn5 + *pilA* (IIB) | *pilA* transposon mutant complemented with pBADGr-*pilA* (IIB) | This work |

| | | |
|---|---|---|
| mPAO1 *pilA*::Tn5 + *pilA* (IIC) | *pilA* transposon mutant complemented with pBADGr-*pilA* (IIC) | This work |
| mPAO1 *pilA*::Tn5 + *pilA-tfpY* | *pilA* transposon mutant complemented with pBADGr-*pilA-tfpY* | (5) |
| mPAO1 *pilA*::Tn5 + *pilA-tfpW-tfpX* + D-*araf* | *pilA* transposon mutant complemented with pBADGr *pilA-tfpW-tfpX* and pUCP20-D-*araf* | (8) |
| mPAO1 *pilA*::Tn5 + *pilA-tfpZ* | *pilA* transposon mutant complemented with pBADGr-*pilA-tfpZ* | (5) |
| PA1244 | Group I clinical isolate | (11) |
| PA14 | Group III lab strain | (12) |
| PA5196 | Group IV clinical isolate | (6) |
| PA1457 | Group V clinical isolate | (6) |
| mPAO1 *pilA*::Tn5 + *pilA* (PA1244) | *pilA* transposon mutant complemented with pBADGr-*pilA* (PA1244) | (6) |
| mPAO1 *pilA*::Tn5 + *pilA* (PA13756) | *pilA* transposon mutant complemented with pBADGr-*pilA* (PA13756) | This work |
| mPAO1 *pilA*::Tn5 + *pilA* (PA13877) | *pilA* transposon mutant complemented with pBADGr-*pilA* (PA13877) | This work |
| mPAO1 *pilA*::Tn5 + *pilA* (PA14868) | *pilA* transposon mutant complemented with pBADGr-*pilA* (PA14868) | This work |
| mPAO1 Δ*pilA* | Deletion of *pilA* | (13) |
| mPAO1 Δ*pilA* + EV | *pilA* mutant complemented with pHERD30T | This work |
| mPAO1 Δ*pilA* + *pilA* | *pilA* mutant complemented with pHERD30T-*pilA* | This work |
| mPAO1 Δ*pilA* + PilA K65E | *pilA* mutant complemented with pHERD30T-PilA K65E | This work |
| mPAO1 Δ*pilA* + PilA K67E | *pilA* mutant complemented with pHERD30T-PilA K67E | This work |
| mPAO1 Δ*pilA* + PilA E77K | *pilA* mutant complemented with pHERD30T-PilA E77K | This work |

| | | |
|---|---|---|
| mPAO1 Δ*pilA* + PilA E83K | *pilA* mutant complemented with pHERD30T-PilA E83K | This work |
| mPAO1 Δ*pilA* + PilA D85K | *pilA* mutant complemented with pHERD30T-PilA D85K | This work |
| mPAO1 Δ*pilA* + PilA E97K | *pilA* mutant complemented with pHERD30T-PilA E97K | This work |
| mPAO1 Δ*pilA* + PilA D98K | *pilA* mutant complemented with pHERD30T-PilA D98K | This work |
| mPAO1 Δ*pilA* + PilA D103K | *pilA* mutant complemented with pHERD30T-PilA D103K | This work |
| mPAO1 Δ*pilA* + PilA K116E | *pilA* mutant complemented with pHERD30T-PilA K116E | This work |
| mPAO1 Δ*pilA* + PilA K120E | *pilA* mutant complemented with pHERD30T-PilA K120E | This work |
| mPAO1 Δ*pilA* + PilA D129K | *pilA* mutant complemented with pHERD30T-PilA D129K | This work |
| mPAO1 Δ*pilA* + PilA K135E | *pilA* mutant complemented with pHERD30T-PilA K135E | This work |
| mPAO1 Δ*pilA* + PilA D139K | *pilA* mutant complemented with pHERD30T- PilA D139K | This work |
| mPAO1 Δ*pilA* + PilA K67E E77K | *pilA* mutant complemented with pHERD30T- PilA K67E E77K | This work |
| Bacteriophages | | |
| Kipling | DsDNA LPS-targeting bacteriophage | This work |
| JBD5 | DsDNA pilus-targeting bacteriophage | (14) |
| JBD18 | DsDNA pilus-targeting bacteriophage | (15) |
| JBD25 | DsDNA pilus-targeting bacteriophage | (15) |
| JBD26 | DsDNA pilus-targeting bacteriophage | (16) |
| JBD68 | DsDNA pilus-targeting bacteriophage | (16) |
| JBD93 | DsDNA pilus-targeting bacteriophage | (16) |

| | | |
|---|---|---|
| DMS3 | DsDNA pilus-targeting bacteriophage | (17) |
| MP22 | DsDNA pilus-targeting bacteriophage | (18) |
| PO4 | DsDNA pilus-targeting bacteriophage | (19) |
| Cootes | DsDNA pilus-targeting bacteriophage | This work |
| Lindberg F10 | DsDNA pilus-targeting bacteriophage | (20) |

**Supplementary Table S4. Primers used in this study.**

| Gene | Forward (5’ → 3’) | Reverse (5’ → 3’) |
|---|---|---|
| *pilA* | TTTTGAATTCGAGAGATTCATGAAAGCTCAAAAAGGCTTTAC | TTTTAAGCTTTTAGTTATCACAACCTTTCGGAGTGAACATC |
| PilA K65E | GAATTGCTGGTAGCGAAATTAAAATTGGTACTAC | GTAGTACCAATTTTAATTTCGCTACCAGCAATTC |
| PilA K67E | CTGGTAGCAAAATTGAAATTGGTACTACTGC | CTGGTAGCAAAATTGAAATTGGTACTACTGC |
| PilA E77K | CTTCTACTGCGACCAAAACATATGTCGG | CCGACATATGTTTTGGTCGCAGTAGAAG |
| PilA E83K | GGCGTCAAACCGGATGCCAA | TTGGCATCCGGTTTGACGCC |
| PilA D85K | GTCGAGCCGAAAGCCAACAAGT | ACTTGTTGGCTTTCGGCTCGAC |
| PilA E97K | CTGTAGCAATCAAAGATAGTGGTGCGG | CCGCACCACTATCTTTGATTGCTACAG |
| PilA D98K | GTAGCAATCGAAAAAAGTGGTGCGG | CCGCACCACTTTTTTCGATTGCTAC |
| PilA D103K | GATAGTGGTGCGGGTAAAATTACCTTTACC | GGTAAAGGTAATTTTACCCGCACCACTATC |
| PilA K116E | CCTCTAGTCCCGAAAATGCTACTAAAGTTATC | GATAACTTTAGTAGCATTTTCGGGACTAGAGG |
| PilA K120E | CCCAAGAATGCTACTGAAGTTATCACTCTG | CAGAGTGATAACTTCAGTAGCATTCTTGGG |

| | | |
|---|---|---|
| PilA D129K | CGTACTGCGAAAGGGGTCTGG | CCAGACCCCTTTCGCAGTACG |
| PilA K135E | TCTGGGCTTGTGAATCTACCCAGG | CCTGGGTAGATTCACAAGCCCAGA |
| PilA D139K | CTTGTAAATCTACCCAGAAACCGATGTTCAC | GTGAACATCGGTTTCTGGGTAGATTTACAAG |
